# Effects of Oxytocin and p-Cresol on BDNF Expression in Pancreatic Cancer Cells

**DOI:** 10.1101/2025.03.28.646013

**Authors:** Gigi Tevzadze, Nino Javakhishvili, Moe Goto, Ivane Abiatari

## Abstract

This study demonstrates for the first time how p-cresol and oxytocin influence BDNF expression in pancreatic cancer cells. Specifically, it was found that while treated with p-cresol, an organic compound produced by gastrointestinal bacteria, BDNF expression in pancreatic cancer cells increases. However, while co-treated with oxytocin, a peptide hormone, BDNF expression decreases in these cells.

## Introduction

Oxytocin is a peptide hormone and neuropeptide produced in the hypothalamus and secreted by the posterior pituitary gland.^1^ It plays a significant role in both human and animal behavior.^2,3^ The release and action of oxytocin are commonly associated with the intensification of social bonds, parent-child relationships, sexual behavior, and childbirth.^3^ Medically, oxytocin is primarily used to enhance uterine contractions during childbirth.^5^

In recent decades, studies have frequently linked oxytocin to autism.^6^ However, a definitive association between intranasal oxytocin administration and behavioral improvement in individuals with autism spectrum disorder has yet to be established. Additionally, in animal studies, intranasal oxytocin administration has been associated with enhanced learning abilities^7^ and delayed onset of Alzheimer’s disease.^8^

Animal studies have also demonstrated that oxytocin improves regenerative processes and slows aging.^9^ Measurements of oxytocin levels in human blood plasma indicate a significant decline with age.^9^

Recent research has examined the correlation between brain-derived neurotrophic factor (BDNF) levels and cancer cell growth. BDNF is a neurotrophic factor implicated in promoting cancer cell growth and proliferation.^10^ Elevated BDNF levels have been observed in several human cancer types, including prostate, lung, and breast cancers, as well as in adjacent tissues, all of which are associated with high mortality rates.^10^

Moreover, emerging studies suggest a potential relationship between oxytocin and BDNF. Research indicates a correlation between oxytocin and BDNF levels,^11^ and previous findings from our group demonstrated a decrease in BDNF release in P12 cells following oxytocin administration.^12^ Particularly, this study demonstrated that oxytocin has a statistically significant effect on P-12 cells (namely, termination of BDNF-mediated growth) in the presence of p-Cresol administration. In the same study, it was observed that the independent administration of p-Cresol increases BDNF levels in P-12 cells.

resol, also known as 4-methylphenol, is an organic compound produced by colonies of gastrointestinal bacteria.^13^ In recent years, various studies have highlighted its association with mental disorders.^14^ Furthermore, p-Cresol has been shown to persist in statistically significant concentrations in human blood, irrespective of age, health issues or diet.^15^ As previously noted, the administration of oxytocin to nerve cells exerted an effect only when combined with p-Cresol. Independently, however, p-Cresol was found to increase BDNF levels in P-12 cells.

Several explanations for this phenomenon are possible; However, this factor should be considered in any *in vitro* experiments involving cell development.

The current experimental design is based on the following data:

1. Experimental evidence suggests that oxytocin decreases BDNF levels in P12 cells or is at least negatively correlated with BDNF increases in co-administration with p-cresol.
2. Elevated BDNF levels are associated with the growth of human cancer cells.
3. Oxytocin release is closely linked to social bonds and interactions, while its levels decrease with age.
4. Cancer incidence statistics reliably indicate an increased likelihood of developing cancer with age.^16^
5. The constant amount of p-Cresol in human blood regardless of age and other factors and its effect on the increase in BDNF in particular cells.

Based on these assumptions, we thought to investigate the effect of p-Cresol and oxytocin on BDNF expression, particularly in pancreatic cancer cells.

## Materials and Methods

### Reagents

The Panc-1, pancreatic cancer cell line was a gift from the Pancreatic Research Lab of the Technical University of Munich, Germany. p-Cresol was purchased from UHN Shanghai R&D Co. Ltd. The human BDNF ELISA Kit (EH42RB), Oxytocin 96% (J63421.LB0), High glucose Dulbecco’s modified Eagle’s medium (DMEM), and Fetal Bovine Serum (FBS) were purchased from Thermo Fisher Scientific.

### Cell Culture and Treatment

Panc-1 pancreatic cancer cells were routinely grown in DMEM supplemented with 10% FCS, 100 units/mL penicillin, and 100 μg/mL streptomycin (standard medium). Cells were maintained at 37°C in a humid chamber with 5% CO_2_ and 95% air atmosphere and regularly tested for mycoplasma contamination. For the experiment, 2 million cells were seeded in 6-well tissue culture plates (Sigma-Aldrich) in 5ml standard medium. After 12 hours, the standard medium was replaced with the treatment medium (DMEM supplemented with 1% FCS, 100 units/mL penicillin, and 100 μg/mL streptomycin). After another 12 hours, the reagent-containing treatment mediums were applied to the cells in four different groups: 1) control (the treatment medium only), 2) p-Cresol (1 µM), 3) Oxytocin (1 µM) and 4) p-Cresol / Oxytocin (1 µM/1 µM) and incubated for 5 days. Medium from each treatment group was collected for the BDNF concentration measurement on the 3^rd^ and 5^th^ days of the treatment and, after brief centrifugation (1,500 rpm/5 min/RT), were frozen at -20°C.

### BDNF assessment

The BDNF concentration was measured in the medium samples using the Human BDNF ELISA Kit, following the manufacturer’s protocol. Briefly, 100 µL of the diluted standards and samples were added to wells and incubated for 2.5 hours at room temperature. Next, the wells were washed, and 100 µL of biotinylated anti-BDNF antibody was added to each well for 1 hour at room temperature with gentle shaking. After washing away the unbound biotinylated antibody, 100 µL of HRP-conjugated streptavidin was pipetted into wells for 45 minutes at room temperature. Then, the wells were washed, and TMB (3,3′,5,5′-tetramethylbenzidine) substrate solution was added to the wells for 30 minutes at room temperature in the dark with gentle shaking. After adding the stop solution, the color intensity was measured at 450 nm using an ELx808 microplate spectrophotometer (BioTek). The BDNF concentration in each sample was calculated based on the standard sample curve generated within the assay.

### Statistical analysis

Data were obtained from three independent culture experiments. Statistical analysis and graph presentation were performed using the GraphPad Prism Software (GraphPad, San Diego, Calif., USA). The results are expressed as the mean ± standard error of the mean. For the comparisons of groups, one-way analysis of variance (ANOVA) for random measures was applied. The level of statistical significance was set at p < 0.05.

## Results

Previous work showed that p-Cresol increases BDNF secretion in PC-12 cells.^12^ In this study, we analyzed whether p-Cresol affected BDNF secretion in Panc-1 cells and whether Oxytocin can mediate p-Cresol-stimulated BDNF expression. Results showed that the addition of p-Cresol increased BDNF secretion in Panc-1 cells after three days (2.33 ± 0.11 ng/ml) and five days (2.43 ± 0.3 ng/ml) of treatment compared to control (1.89 ± 0.05 and 2.25 ± 0.02 ng/ml respectively). Interestingly, oxytocin co-treatment significantly reduced p-Cresol-mediated BDNF secretion in Panc-1 cells with a concentration of 1.17 ± 0.17 ng/ml after three days and 1.8 ± 0.15 ng/ml after five days of treatment. Oxytocin treatment alone did not induce marked changes in BDNF expression compared to controls (Figure 1).

**Figure 1.**
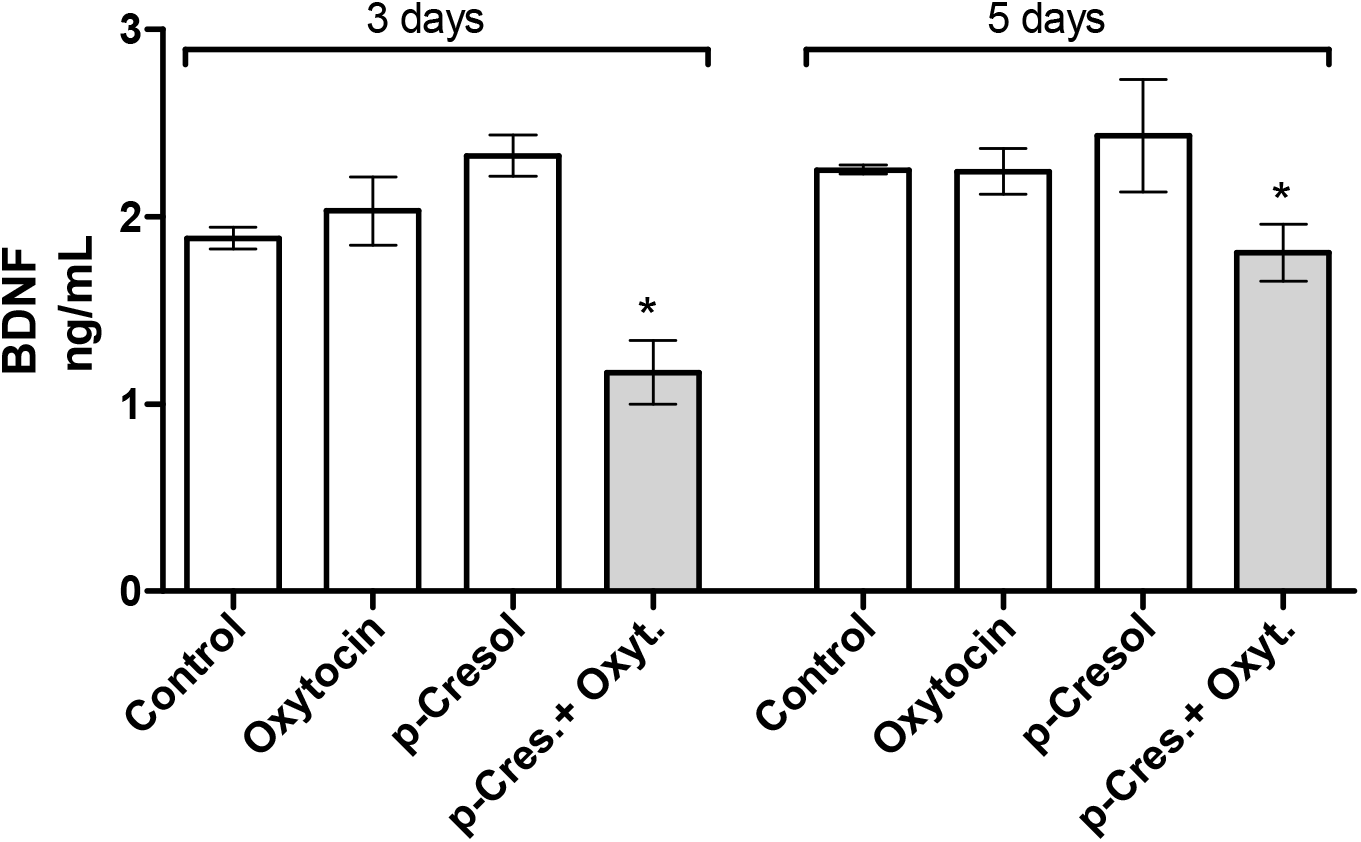
Analysis of the BDNF expression by Panc-1 pancreatic cancer cells. ELISA demonstrated concentration of BDNF in cell supernatants after 3 and 5 days of 1 µM p-Cresol and 1 µM Oxytocin treatment/co-treatment. Data is presented as mean ± SEM of three independent experiments. *\*p < 0*.*05*.

## Discussion

Our study demonstrated that a specific dose of oxytocin can significantly influence the BDNF levels in cancer cells. Specifically, this effect was observed *in vitro*, where oxytocin inhibited the p-Cresol-mediated production of BDNF in pancreatic cancer cells.

Our study also demonstrated that the separate administration of p-Cresol led to increased BDNF levels in these cells. These findings suggest a potential connection between the effects of oxytocin and p-Cresol on BDNF regulation in cancer cells. Specifically, the observed decrease in BDNF levels upon oxytocin administration and the increase in BDNF levels with p-Cresol may indicate systemic, organism-internal factors contributing to cancer cell growth and development.

This finding aligns with two key statistical observations: the increased probability of cancer development with age^16^ and the significant decline in oxytocin secretion over time.^9^ Based on these results, it can be hypothesized that oxytocin and p-Cresol may play a crucial role in the body, particularly in oncogenesis.

Further research is necessary to test this hypothesis comprehensively. Future studies should include both *in vitro* and *in vivo* models to investigate not only changes in BDNF levels but also the direct impact of oxytocin and p-Cresol on the pathophysiology of cancer cells.

## Conclusion

This study provides the first evidence that BDNF levels are increased in pancreatic cancer cells following p-Cresol treatment and reduced upon Oxytocin addition. Given the proposed role of BDNF in promoting the growth of certain cancer types, these findings may open new avenues for both research and potential therapeutic applications.

